# Doublecotin-like kinase 1 increases chemoresistance of colorectal cancer cells through the anti-apoptosis pathway

**DOI:** 10.1101/517425

**Authors:** Lianna Li, Kierra Jones, Hao Mei

## Abstract

Colorectal cancer (CRC) is the third most common cancer diagnosed and the second leading cause of cancer-related deaths in the United States. About 50% of CRC patients relapsed after surgical resection and ultimately died of metastatic disease. Cancer stem cells (CSCs) are believed to be the primary reason for the recurrence of CRC. Specific stem cell marker, doublecortin-like kinase 1 (DCLK1) plays critical roles in initiating tumorigenesis, facilitating tumor progression, and promoting metastasis of CRC. It is up-regulated in CRC and up-regulation of DCLK1 indicates poor prognosis. Whether DCLK1 is correlated with enhanced chemoresistance of CRC cells is unclear. Our research aims to reveal association of DCLK1 with chemoresistance of CRC cells and the underlying molecular mechanisms. In order to achieve our goal, we established stable DCLK1 over-expression cells (DCLK1+) using the HCT116 cells (WT). DCLK1+ and WT cells were treated with 5-Fluorouracil (5-Fu) at different doses for 24 or 48 hours. MTT assay was used to evaluate cell viability and IC_50_ of 5-Fu was determined. Quantitative real time PCR was applied to determine gene expression of caspase-3 (casp-3), caspase-4 (casp-4), and caspase-10 (casp-10). Cleaved casp-3 expression was investigated using Western blot and immunofluorescence. Our results demonstrated that IC_50_ of 5-Fu for the DCLK1+ cells was significantly higher than that of the WT cells for both 24 and 48-hour treatment (*P*=0.002 and 0.048 respectively), indicating increased chemoresistance of the DCLK1+ cells. Gene expression of casp-3, casp-4, and casp-10 were significantly inhibited in the DCLK1+ cells after 5-Fu treatment compared to the WT cells (*P*=7.616e-08, 1.575e-05 and 5.307e-08, respectively). Cleaved casp-3 amount and casp-3 positive cells were significantly decreased in the DCLK1+ cells after 5-Fu treatment compared to the WT cells (*P*=0.015). In conclusion, our results demonstrated that DCLK1 overexpression enhanced the chemoresistance of CRC cells to 5-Fu treatment by suppressing gene expression of key caspases in the apoptosis pathway and activation of apoptosis pathway. DCLK1 can be an intriguing therapeutic target for the effective treatment of CRC patients.

## Introduction

Colorectal cancer (CRC) is the third most common cancer diagnosed in both men and women and the second leading cause of cancer-related deaths in the United States (http://www.cdc.gov/cancer/colorectal/statistics/). In the last 20 years, progress in the treatment of CRC has improved quality of patients’ life, but up to 50% of patients relapsed after surgical resection and ultimately died of metastatic disease [1]. Adjuvant systemic chemotherapy with cytotoxic drugs is recommended as standard clinical practice for patients with stage III CRC after surgical resection of the local CRC [2] since the survival outcomes of CRC patients with adjuvant systemic chemotherapy combined with surgical resection was significantly higher than those with surgical resection only [3]. The promising progress of systemic chemotherapy for CRC began with the discovery of 5-fluorouracil (5-Fu) in 1957 [4]. Currently the conventional first-line treatments for CRC patients are the combination of 5-Fu, leucovorin, and oxaliplatin (FOLFOX) or the combination of 5-Fu, leucovorin, and irinotecan(FOLFIRI) [5]. However, not all of the CRC patients respond to the systemic therapies, and even though for the responsive patients, almost all of them developed resistance [6]. According to the cancer stem cell (CSC) hypothesis, presence of chemoresistant CSCs (also known as tumor stem cells (TSCs)) is the primary cause [7, 8]. CSCs accounts for 0.05-1% of the tumor mass, but they can give rise to all of the cell types in the tumor and possess unlimited self-renewal capability [9]. Several specific putative markers have been identified for the stem cell populations in the gastrointestinal tract, including doublecortin-like kinase 1 (DCLK1, also known as KIAA0369 [10] or DCAMKL1[11]).

DCLK1 is a microtubule-associated serine-threonine protein kinase and functions in facilitating polymerization of tubulin dimers to assemble microtubules [12]. It is predominantly expressed in the nervous system and is correlated with normal nervous system development [13, 14] and general cognition and verbal memory function [15]. In the late 2000s, DCLK1 was identified as a stem cell marker for the intestinal stem cells and correlated with stemness of CRC cells [11, 16]. It is co-localized with other well-characterized gastrointestinal stem cell markers, such as Lgr5 in the “+4 position” of the crypt of small intestine where the intestinal stem cells are located [17, 18]. Up-regulated expression of DCLK1 was found broadly in solid tumors almost all over the body, including the esophageal cancer [19], pancreatic cancer [20, 21], liver cancer [22], CRC [23, 24], etc. and is correlated with poor prognosis [25]. The most recent clinical findings identified that elevated DCLK1+ cells in the blood can be used as a novel non-invasive marker for the diagnosis of incidence, relapse and metastasis for CRC [26], liver cancer [27], pancreatic cancer [28] and Barrett’s esophagus and esophageal adenocarcinoma [29]. DCLK1 played important roles in the initiation, progression and metastasis of CRC [30-32]. It can promote cell survival via the prevention of cancer cell apoptosis in neuroblastoma [33] and anoikis in mouse colonic epithelial cells [34].

Though DCLK1 is such a multiple functional protein in the CRC tumorigenesis, neither association of DCLK1 with chemoresistance in human CRC nor the underlying cellular and molecular mechanism is clear. In this paper, we identified that DCLK1 can significantly increase chemoresistance of CRC cells to 5-Fu treatment, and it functions through inhibition of gene expression of key caspases and activation of apoptosis pathway. Our results demonstrated that DCLK1 can be used as an intriguing therapeutic target for CRC treatment.

## Materials and Methods

### Cell line and cell culture

Human colorectal carcinoma cell line HCT116 cells were purchased from ATCC (ATCC® CCL-247™) and were maintained in McCoy’s 5A medium (ATCC® 30-2007™) supplemented with 10% FBS in 37 °C incubator with 5% CO2. Isogenic DCLK1 over-expressed cells (DCLK1+) were established by transfecting human DCLK1 variant 1 cDNA, which is fused with a turboGFP gene at C-terminal (OriGene, Cat # RG217050) into HCT116 cells. In order to avoid the clonal variance, different DCLK1 over-expressed clones were selected. Control HCT116 cells (WT) were established by transfecting pCMV6-AC-GFP Tagged Cloning Vector (Origene, Cat # PS100010) into HCT116 cells. Both DCLK1 over-expressed cells and control HCT116 cells were selected (400ug/ml) and maintained (250ug/ml) using Geneticin(G418).

### 5-Fu cytotoxicity assay

WT and DCLK1+ cells were plated at 1×10^4 cells/well/100µL in the 96-well plate for 24 hours. Then cells were treated with 5-Fu (Sigma; F6627-1G) at different concentrations for 24 or 48 hours. Cell viability was determined by MTT assay according to Li’s approach with modifications [35]. Briefly, MTT reagent (5mg/ml) was added into cells at a 1:10 ratio of the culture medium and incubated for 3 hours at 37°C. After incubation, culture medium with MTT was replaced by dimethyl sulfoxide (DMSO). Plate was sent to the BioTek Synergy 2 multi-mode reader and absorbance was measured at 570nm and 630nm. OD value used to establish the dose-killing curve was achieved by subtracting OD630 (background) from OD570. IC_50_ of 5-Fu was calculated using equation from the established dose-killing curve by setting the cell viability at 50%.

### Western Blotting

WT and DCLK1+ cells were plated at 2 x 10^6 cells per T-25 flask and cultured for 24 hours. Then cells were treated with/without 5-Fu and cultured for 48 more hours. Whole cell lysates were harvested using ice-cold RIPA buffer with 1x protease inhibitor (Sigma, P8340) and 1x phosphate inhibitor (Sigma, P5726). Protein concentration was determined using Pierce™ BCA Protein Assay Kit according to the manufacture’s manual (ThermoFisher Scientific, Catalog number: 23227). 40µg of protein from each sample was loaded onto the 4– 15% Mini-PROTEAN^®^ TGX™ Gel (BioRad, Catalog number: 4561083) for sodium dodecyl sulfate-polyacrylamide gel electrophoresis (SDS-PAGE) at 100V for 60 minutes. Proteins were transferred to nitrocellulose membrane using the Trans-Blot Turbo System (Bio-Rad). Then the membrane was blocked in 3% non-fat milk in 1X PBS with 0.1% tween 20 (1xPBST) at room temperature with constant shaking for one hour and was probed with primary antibodies (caspase-3 (Life Biosciences, LS-C331947) and cleaved caspase-3 (Life Biosciences, LS-C380472)) in 1X PBST with 3% non-fat milk at 4°C overnight. After the primary antibody incubation, membrane was washed with 1xPBST for 5 minutes, 3 times total. Then the membrane was probed with secondary antibody with 1% non-fat milk at room temperature for an hour. Membrane was washed with 1xPBST for 5 minutes, 4 times total, and sent for imaging using the LI-COR Odyssey Imaging System.

### Immunofluorescence

WT and DCLK1+ cells were plated at 3 x 10^5 cells/well into 8-well Millicell EZ chamber slides (Millipore, cat. no. PEZGS0816) overnight. Cells were treated with/without 5-Fu for 24 hours and cellular cleaved caspase-3 protein was labeled immunofluorescently. Briefly, culture medium was removed and cells were washed 3x with 1x PBS. Then cells were fixed with 4% paraformaldehyde for 10 minutes on ice. Wash cells 3x with 1x PBS, and block with 1% BSA by incubate cells for 5 min at 40°C. Wash cells 3x with 1x PBS and incubate cells with the primary antibody in 1x PBS with 1% BSA overnight at 4°C. Wash cells 5x with 1x PBS and probed with secondary antibody by incubating for 1 hour at room temperature in dark. Wash cells 5x with 1x PBS, and did nuclei counterstain using 1X DAPI (Sigma cat. no. D9542) by incubating for 15 min at room temp. Wash cells with ddH2O, remove gasket, mounting with Prolong Gold Anti-fade reagent (#9071) and cover with coverslip. Images were taken using a Life Technologies EVOS FL fluorescent microscope.

### Quantitative real time polymerase chain reaction [(q)RT-PCR]

Total cellular RNA was isolated using the RNeasy Mini Kit (Qiagen, Valencia, CA) according to the manufacturer’s instructions from the WT and DCLK1+ cells. First strand cDNA was generated using the Reverse Transcription System (Promega, Madison, WI) according to the manufacturer’s instruction. 5 µL cDNA from reverse transcription PCR was added to a 25 µL reaction containing sybergreen. Primers for the human GAPDH, caspase 3, 4 and 10 are listed in Table 1. (q)RT-PCR was carried out on a Stratagene Mx3005 quantitative real time PCR thermocycler according to the manufacturer’s instruction. Expression of gene of interests was normalized to GAPDH first, then comparison of fold change between DCLK1+ and WT cells was carried out using the 2^-ΔΔCt^ approach [36].

**Table 1.**
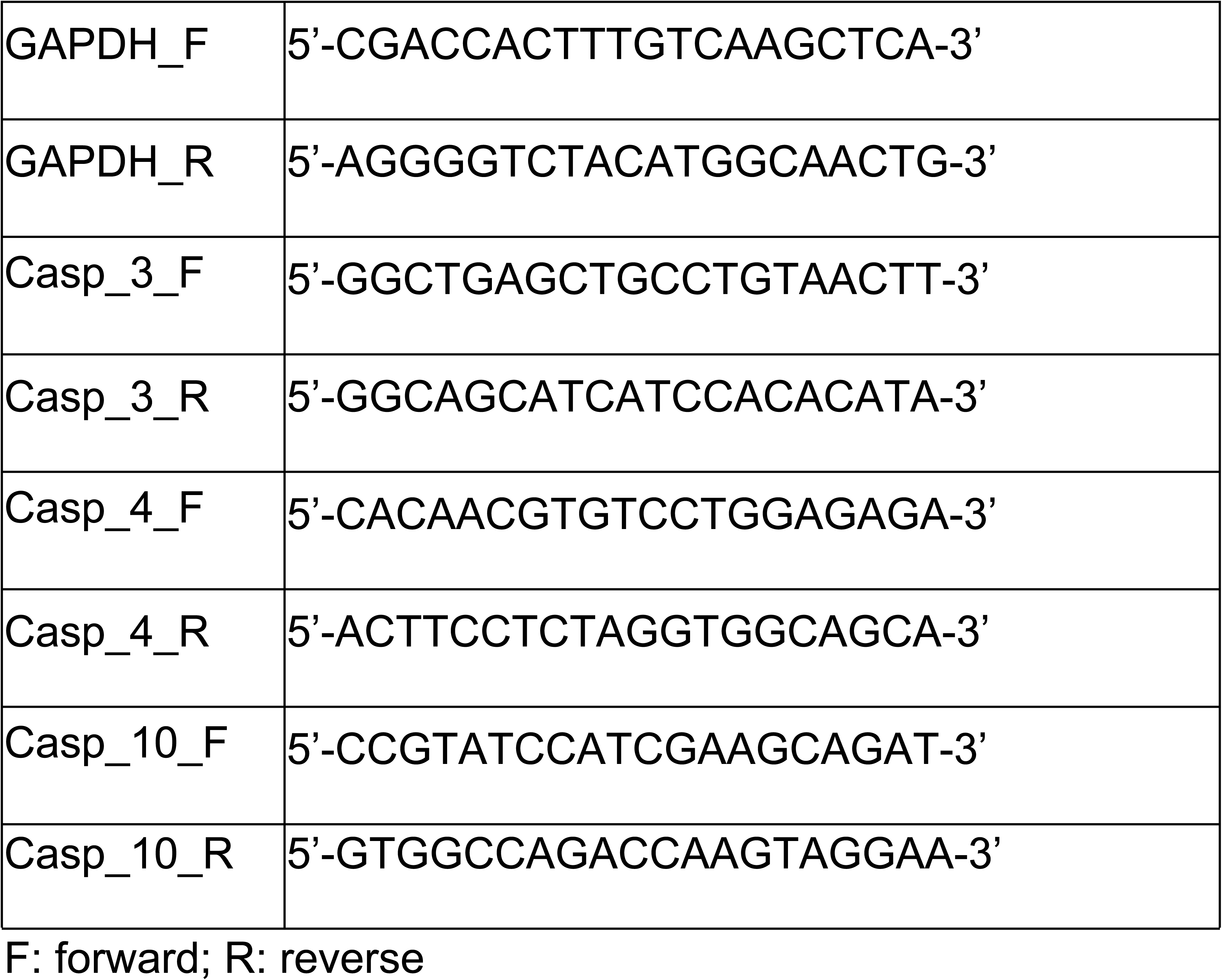
Primers used for the quantitative real-time PCR.

### Statistical analysis

All data was expressed as 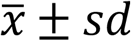. Non-paired one-side *t*-test, two-side *t*-test or paired two-side *t*-test was applied. *P*<0.05 was considered as statistical significant.

## Results

### DCLK1 increases chemoresistance of colorectal cancer cells

In order to investigate whether DCLK1 modifies sensitivity of CRC cells to chemotherapy, the WT and DCLK1+ cells were treated with different doses of 5-Fu for 24 and 48 hours. Cell viability was measured using MTT assay after the designed time period of drug treatment. Our results demonstrated that as 5-Fu concentration increased, cell viability of the WT and DCLK1+ cells decreased for both 24 and 48-hour treatment. Cell viability of DCLK1+ was higher than the WT cells at every dose for both 24 and 48-hour treatment (Figure 1A). IC_50_ of 5-Fu for the DCLK1+ cells was significantly higher than that of the WT cells for both 24 and 48-hour treatment (*P*=0.002 and 0.048 respectively, Figure 1B), indicating increased chemoresistance of the DCLK1+ cells.

**Figure 1.**
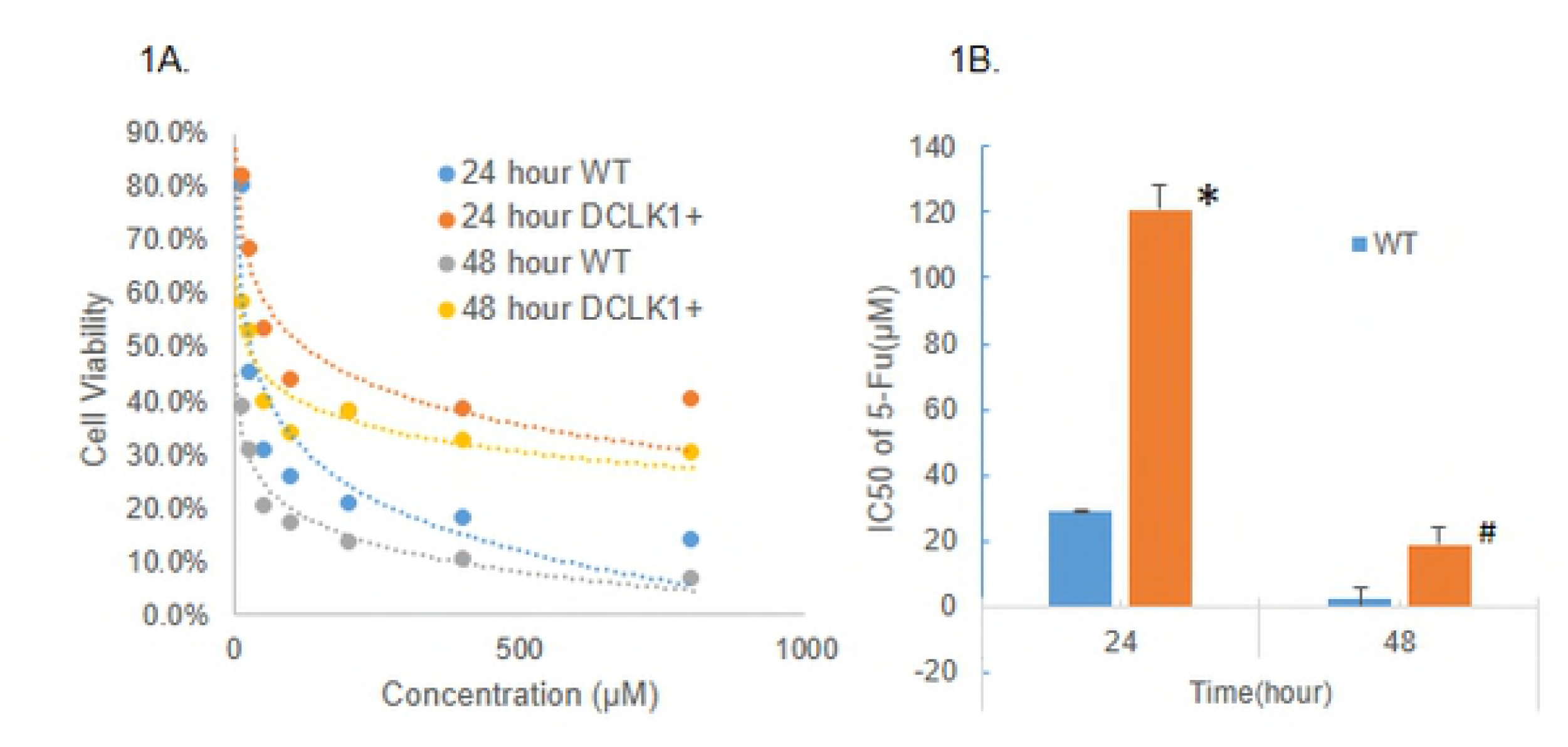
Chemosensitivity of HCT116 wild type cells (WT) and DCLK1 over-expressed cells (DCLK1+) to 5-Fu treatment. **1A.** Cell viability after 5-Fu treatment with different doses at different time from one representative experiment. **1B.** IC50 of 5-Fu for WT and DCLK1+ cells treated for different times. Data was expressed as mean ± SEM from 3 independent experiments. When compared IC50 of 5-Fu in DCLK1+ cells to WT cells, \**P*=0.002 for 24-hour treatment and ^#^*P*=0.048 for 48-hour treatment.

### DCLK1 suppresses gene expression of multiple critical caspases transcriptionally

DCLK1 might be correlated with apoptosis pathway since knockdown of DCLK1 gene expression induced apoptosis of neuroblastoma cells [33] and anoikis-resistant mouse colonic epithelial cells expressed increased amount of DCLK1 [34]. Our RNA-Seq data indicated that DCLK1 over expression modulates gene expression of multiple key caspases, specifically casp-4 and 10 (Figure 2A), which might due to antibiotics G418 selection. In order to find out whether DCLK1 can affect gene expression of casp-4 and 10 without G418 selection but with 5-Fu treatment, and whether casp-3 is also controlled by DCLK1 transcriptionally, (q)RT-PCR was carried out. Our results demonstrated that with G418 selection but without 5-Fu treatment, gene expression of casp-3 and 4 was very similar in the WT and the DCLK1+ cells. Casp-10 was significantly lower in the DCLK1+ cells, which is consistent with the RNA-Seq data. However, in the cells without G418 selection but with 5-Fu treatment, gene expression of all three genes was significantly inhibited in the DCLK1+ cells compared to the WT cells (*P*=7.616e-08, 1.575e-05 and 5.307e-08, respectively, Figure 2B) and casp-10 was further inhibited compared to G418 selection, indicating DCLK1 can significantly inhibit gene expression of casp-3, 4 and 10 transcriptionally, which may be responsible for the higher cell survival rate after 5-Fu treatment and contribute to the increased chemoresistance of the DCLK1+ cells.

**Figure 2.**
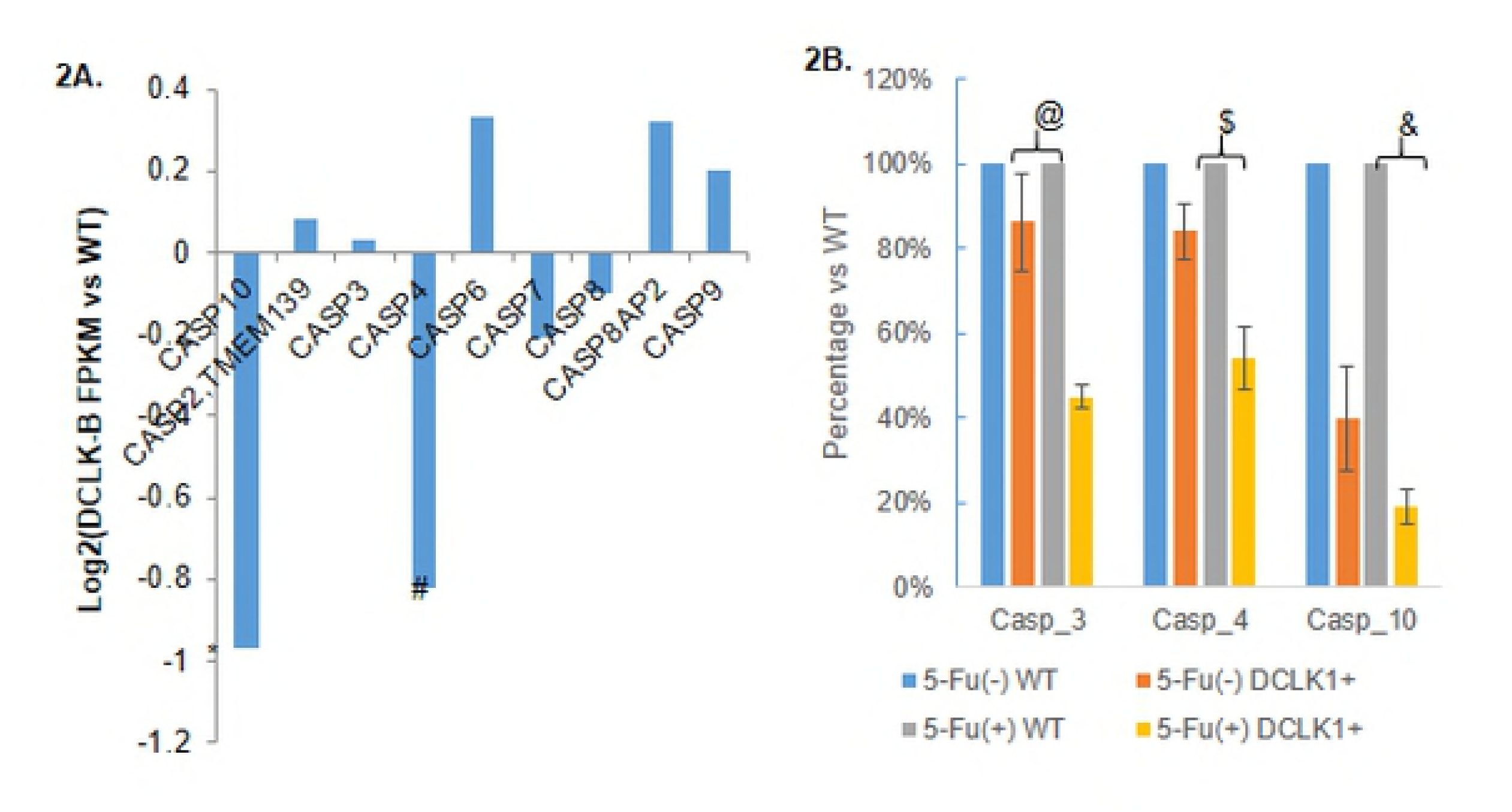
Gene expression of multiple caspases in HCT116 wild type (WT) and DCLK1 over-expressed (DCLK1+) cells. **2A.** Comparison of gene expression of multiple caspases in the WT and DCLK1+ cells without 5-Fu treatment determined by RNA-Sequencing. \**P*=6.68E-04; ^#^*P*=4.16E-04. **2B.** Gene expression of casp-3, 4 and 10 in the WT and DCLK1+ cells with/without 5-Fu treatment determined by quantitative real time PCR. Gene expression of casp-3, 4 and 10 was normalized to GAPDH first before comparison between WT and DCLK1+ cells. Data was expressed as mean ± SEM from 3 independent experiments. ^@^*P=*7.616e-08, ^$^*P*=1.575e-05 and ^&^*P*=5.307e-08.

### DCLK1 suppresses activation of the apoptosis pathway

In order to find out whether DCLK1 also modulate activation of the apoptosis pathway, we evaluated the cleaved casp-3 protein expression level in the WT and the DCLK1+ cells before and after 5-Fu treatment for 24 hours. Our results demonstrated that before 5-Fu treatment, both WT and the DCLK1+ cells demonstrated undetectable cleaved casp-3 expression, indicating no or low apoptosis events. However, with 5-Fu treatment, the WT cells showed a much higher cleaved casp-3 expression level compared to the DCLK1+ cells (Figure 3A), and results of semi-quantification using Image J software demonstrated that amount of cleaved casp-3 is 3 ± 0.8 fold in the WT cells compared to the DCLK1+ cells, indicating much higher apoptosis rate in the WT cells after 5-Fu treatment. When compared the total casp-3 expression level in the WT and DCLK1+ cells, there is no significant difference either with or without 5-Fu treatment.

**Figure 3.**
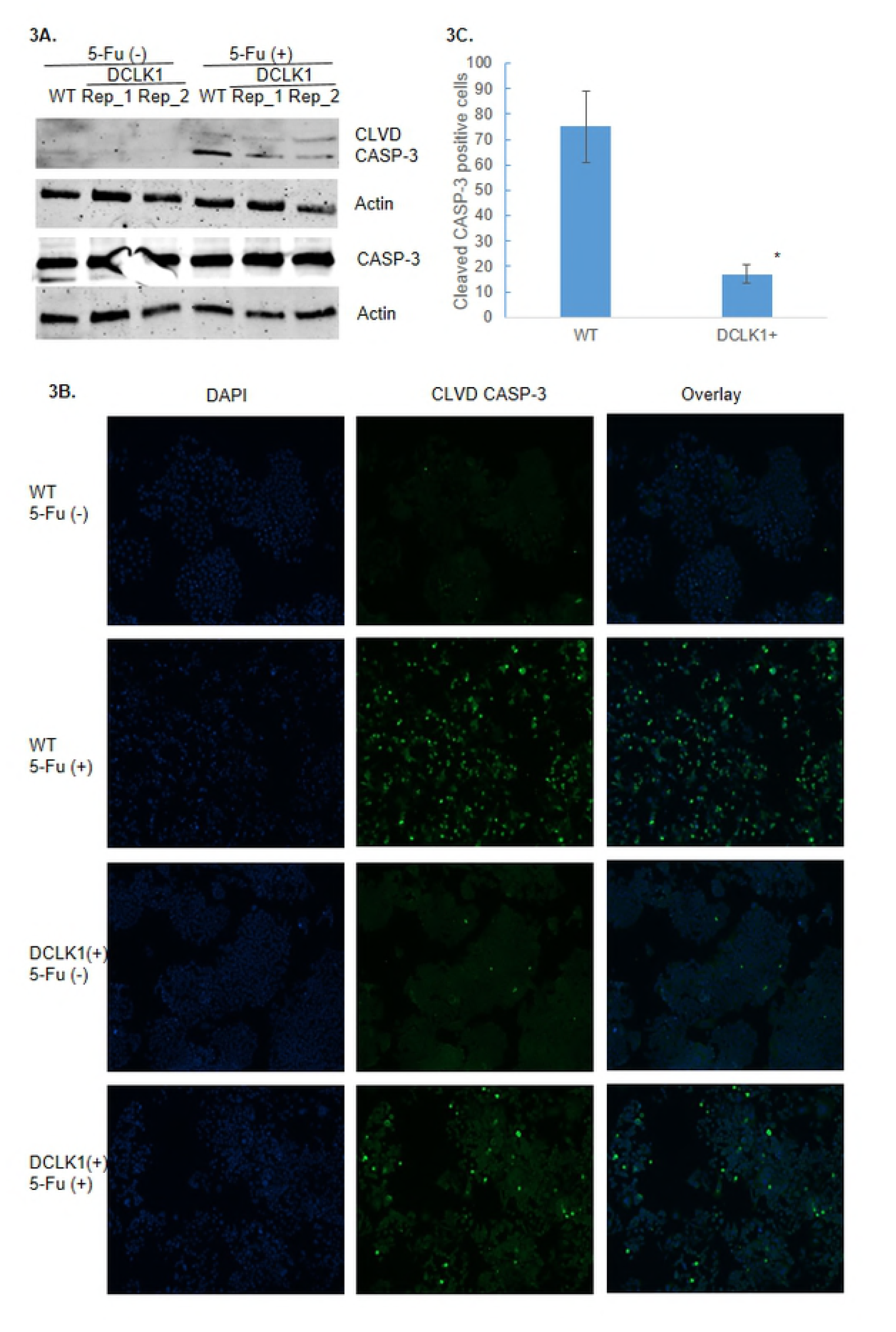
Modulation of activation of apoptosis pathway by DCLK1 in the colorectal cancer cells. **3A.** Expression of cleaved casp-3 (CLVD CASP-3) in the HCT116 wild type (WT) and DCLK1 over-expressed (DCLK1+_Rep_1 and Rep_2) cells with or without 5-Fu treatment determined by Western Blot. Total casp-3(CASP-3) and actin were used as controls. **3B.** Immunofluorescence of CLVD CASP-3 in the WT and DCLK1+ cells with/without 5-Fu treatment. **3C.** Comparison of CLVD CASP-3 positive cells in the WT and DCLK1+ cells after 5-Fu treatment. Data was expressed as mean ± SEM from 5 independent image fields for each cell type. \**P*=0.015.

We further confirmed the Western blot findings with immunofluorescence label of cleaved casp-3 protein inside the WT and DCLK1+ cells after 5-Fu treatment. We found significantly more cleaved casp-3 positive cells in the WT cells than the DCLK1+ cells (*P*=0.015, Figure 3B and 3C). In summary, our data indicated that DCLK1 significantly inhibits activation of the apoptosis pathway post-translationally as well.

### Conclusion and Discussion

Using the DCLK1 over-expressed cells, we identified that DCLK1 can enhance chemoresistance of CRC cells to the 5-Fu treatment significantly through the anti-apoptosis pathway by inhibiting gene expression of critical caspases transcriptionally and post-translationally. This is the first time that the association of DCLK1 with chemoresistance in the CRC cells and the underlying molecular mechaism were clarified.

Chemoresistance is a big challenge for the effective treatment of colorectal cancer. CRC cells have evolved multiple mechanisms to resist and escape from the anticancer drugs treatment. Hypoxia tumor microenvironment [37] and enhancement of autophagy procedure [38] both contribute to the chemoresistance of CRC cells. CRCs can also manipulate expression of multiple microRNAs to facilitate their chemoresistance, including suppressing expression of anti-tumor microRNAs, such as miR-181a/135a/302c [39], miR-874-3p [40], miR-200c [41]; and increasing tumorigenic miRNAs, such as miR-196b-5p [42], miR-315b [43], miR-10b [44]. Several cellular pathways were identified to abnormally modulated to facilitate CRC cells chemoresistant procedure, including activation of Wnt/β-catenin pathway [45], Akt pathway [46], and Notch signalling pathway [47], and inactivation of Hippo signalling pathway [40] and caspase-dependent apoptosis pathway [48]. Recently, it was identified that DCLK1 might be associated with chemoresistance of cancer cells. miR-539, a tumor suppressor microRNA, directly inhibits DCLK1 expression to enhance chemosensitivity of non-small cell lung cancer to cisplatin [49]. Over-expression of DCLK1 significantly increased chemoresistance of kidney cancer cells to the receptor tyrosine kinase inhibitors and mTOR inhibitors [50]. miR-15a might overcome chemoresistance of CRC stem cells to 5-FU through suppression of DCLK1 [51]. However, how DCLK1 enhances chemoresistance of CRC cells is unclear.

In this paper, we identified that DCLK1 can increase chemoresistance of CRC cells through modulate apoptosis pathway. Firstly, it can significantly suppress gene expression of several critical caspases, including casp-3, 4 and 10, after 5-Fu treatment. Secondly, it can significantly inhibit activation of the apoptosis pathway, demonstrated by the decreased expression of cleaved casp-3 after 5-Fu treatment in the DCLK1+ cells. We are not the first one to find that DCLK1 was involved in the apoptosis pathway. It was identified that DCLK1 is essential for the survival of neuroblastoma cells since knockdown of DCLK1 gene expression induced apoptosis of the cells [33] and in the anoikis-resistant mouse colonic epithelial cells, DCLK1 was up-regulated [34]. However, it is the first time that DCLK1 was demonstrated to increase chemoresistance of CRC cells via anti-apoptosis pathway.

DCLK1 plays important roles in the initiation, progression and metastasis of CRC, and targeting DCLK1 is efficient in the inhibition of developed tumor growth in animal model. Using Apc^min/+^ mouse model, Chandrakesan and colleagues demonstrated that DCLK1 was over-expressed in the small intestine of the elderly mice compared to normal control mice, and the over-expression of DCLK1 facilitated epithelial mesenchymal transition (EMT), which in turn resulted in the colorectal tumorigenesis [30]. Most recently, they identified that DCLK1 over-expression is correlated with enhanced pluripotency and self-renewal capability of intestinal epithelial cells [52]. When DCLK1 was conditionally over-expressed by crossing Dclk1^*creERT2*^ Rosa26R mice to the Apc^min/+^ mice, the over-expression of DCLK1 significantly increased the incidence of intestinal polyps compared to the normal control mice [31]. DCLK1+ tuft cells are responsible for the tumorigenesis of colon cancer in the DCLK1-CreERT transgenic mice [32] and over-expression of DCLK1 in human pancreatic cancer stem cells facilitated the tumor invasion and metastasis [53]. When DCLK1 expression was specifically knockdown in mouse model, fewer polyps and decreased dysplasia were observed [30] and when DCLK1+ cells were specifically targeted in the developed polyps, the CSCs died and the established polyps were rapidly collapsed [31]. All of these findings indicate that DCLK1 can become a promising therapeutic target for the CRC treatment.

In summary, DCLK1 was identified to be associated with increased chemoresistance in the CRC cells, and it function through the anti-apoptosis pathway. It can become a new novel therapeutic target for the efficient treatment of CRC patients. Further *in vivo* studies need to be carried out to determine the association of DCLK1 with chemoresistance and its underlying molecular mechanisms, which will provide more evidence for its clinical application and finally become beneficial to CRC patients treatment.

## Acknowledgement

This work was supported by the Mississippi INBRE, funded by an Institutional Development Award (IDeA) from the National Institute of General Medical Sciences of the National Institutes of Health under grant number P20GM103476. The content is solely the responsibility of the authors and does not necessarily represent the official views of the National Institutes of General Medical Sciences or the National Institutes of Health.

